# Ultrafast light sensing mediated by dermal iridophores in the silverside fish *Hypoatherina tsurugae*

**DOI:** 10.64898/2026.07.12.738009

**Authors:** Masakazu Iwasaka

## Abstract

Light-induced body color changes and direct optical sensing in the skin of amphibians and fishes have been debated for more than half a century. Previously reported chromatophore-mediated photoresponses generally occur on timescales of tens of seconds to minutes and are commonly attributed to intracellular molecular mechanisms, including opsin-dependent signaling pathways. In several invertebrates and amphibians, dermal photoreception has been detected electrophysiologically as rapid neural outputs. Here, I report an ultrafast, nonvisual photoresponse mediated by dermal iridophores in the silverside fish *Hypoatherina tsurugae* (Atherinidae). Upon exposure of living skin to white LED illumination, the light reflection from iridophores was rapidly quenched within seconds. Spectral analysis revealed that this quenching response was most sensitive to blue light compared with green and red illumination. The reflected light recovered to its original twinkling state within approximately 10 s after cessation of illumination. The speed and reversibility of this response suggest that mechanisms beyond slow intracellular structural rearrangements are involved and raise the possibility of neural modulation in addition to intrinsic photoreceptive processes. These findings uncover an unrecognized mode of dermal light sensing and provide insight into bioinspired design principles for artificial optical sensing skins.

## 1. Introduction

Amphibians [1] and fishes [2–5] are known to change their skin coloration in response to environmental light conditions. Several studies have reported evidence of eye-independent, direct optical sensing in the skin of these animals. In particular, opsin proteins have been identified as photosensitive molecules not only in retinal tissues but also in the skin of fishes [6–10]. Octopuses have also been shown to sense light through their skin, despite the absence of eye-like image-forming structures [4, 8]. Both studies demonstrated that octopuses respond specifically to blue light. Furthermore, a study investigating chromatophore expansion under light stimulation reported that blue-light sensitivity in the skin matched the spectral sensitivity of opsins in the eyes of the same species [8]. Historically, body coloration in teleost fishes has been extensively investigated in relation to environmental light conditions, such as light and dark environments [1–6, 8, 9, 11]. The speed of color change in fishes and amphibians has also been examined in previous studies. Among chromatophore cell types, iridophores, erythrophores, melanophores, and xanthophores have been reported to contribute to light-induced color changes through distinct mechanisms [8–22]. Erythrophores and xanthophores function as red and yellow color filters, respectively, whereas melanophores act primarily as light absorbers. In contrast, iridophores regulate light reflection intensity within dermal tissues. Because iridophores contain densely packed guanine platelets [23–25], their periodic internal structures generate structural coloration. In most cases, the chromatophores investigated exhibited apparent color-change responses on the timescale of several minutes to tens of minutes. Studies on the iridophores of neon tetra have demonstrated light sensitivity that alters the reflected light spectrum [5, 6, 14–16]. As a possible mechanism, photosensitive molecules have been reported to exist within iridophore cells [6]. The photoresponse of these cells, involving structural color modification, has been attributed to changes in the orientation of guanine platelets inside the cells [15, 16]. Unlike other chromatophores, iridophores do not contain pigment granules that aggregate or disperse. It has therefore been hypothesized that iridophores utilize guanine platelets to generate optical sensing outputs rather than relying on pigment-based mechanisms. Consistent with this hypothesis, changes in the structural color spectrum were observed to occur over several minutes upon exposure to light stimuli [6, 15, 16].

In recent decades, tilapia erythrophores have been studied to examine the migration of pigment particles, and their photosensitivity has been confirmed [9, 15–17]. The involvement of opsin molecules in light-induced aggregation and dispersion of pigment particles has also been suggested in tilapia erythrophores [9]. Light-induced pigment aggregation has been reported in xanthophores as well [10, 18, 19]. These phenomena have been observed in three species, and the involvement of opsins has been discussed in relation to specific responses to blue light. Aggregation and dispersion of melanin particles in melanophores in response to background light intensity have been reported for a long time [3, 4, 20–22]. In melanophores, xanthophores, and erythrophores, light responses generally appear as aggregation or dispersion of pigment particles. In contrast, the photoresponse of iridophores is characterized by changes in the inclination of light-reflecting platelets [5, 12–14]. Many studies have attributed the mechanisms underlying photoresponses to intracellular molecules such as opsins. Similarly, in iridophores, the possible existence and involvement of intracellular opsin molecules have been suggested [6, 7–10]. As mentioned above, the response speed of changes in the light-reflection spectrum of iridophores has been reported to occur on a timescale of minutes, comparable to pigment aggregation and dispersion responses in other chromatophores. In contrast, a recent electrophysiological study in octopus revealed neural responses to light stimulation on the order of milliseconds.

This work presents evidence of a very rapid photoresponse occurring in iridophores of *Hypoatherina tsurugae*, a silverside fish species belonging to the family Atherinidae. The photoresponses observed in the skin of this species were visualized in submillimeter regions of skin tissue containing chromatophores. The light-sensing signal manifests as changes in the light-reflecting behavior of the iridophores in *Hypoatherina tsurugae*.

## 2. Materials and methods

### 2.1 Animals

Specimens of *Hypoatherina tsurugae* (cobalt silverside) were collected by fishing in Fukuoka and Hiroshima, Japan, during June 2022 and October 2024. The fish were maintained in aquaria containing seawater throughout the experiments. A total of 22 specimens were examined in this study. The study was approved by the institutional animal care and use committee (2F19–2, Hiroshima University) and carried out in accordance with all national guidelines including the guidelines for proper conduct of animal experiments (Science Council of Japan). The mean body length and body weight of *H. tsurugae* were 112.1 ± 7.6 mm and 15.4 ± 2.4 g, respectively. In vivo observations of light-reflecting dermal tissues were conducted following the methods described in previous studies [26, 27]. For optical measurements, anesthetized fish were placed in an aquarium. Anesthesia was maintained using tricaine methanesulfonate (50–100 mg L^−1^; Tokyo Chemical Industry, Japan) and 2-phenoxyethanol (0.1%; Fujifilm Wako Pure Chemical Co., Japan).

### 2.2 Image capture of dermal light reflection

Light reflection from the fish body surface was observed using a mono-zoom lens and side illumination (Supplementary Fig. S1). The imaging lens and illumination light source were arranged independently. The experimental setup consisted of a mono-zoom microscope lens (NAVITAR 2.0 × 1-51473, Navitar, Rochester, USA), a CMOS camera (HOZAN L-835, HOZAN, Osaka, Japan), and a white LED light source (LA-HDF158A, Hayashi Repic Co. Ltd., Tokyo, Japan).

Time-series images, including both still images and videos, were captured by the CMOS camera. The white LED illumination used for imaging was aligned such that the central axis of the light beam was parallel to the anteroposterior axis of the specimen. The tip of the light-emitting guide was positioned approximately 30 mm from the parietal region of the fish (Fig. S1). The incident angle of the illumination beam relative to the water surface covering the dorsal trunk ranged from 47° to 56°. Video recordings of microscopic images were acquired using video capture software (Xploview, VIXEN, Saitama, Japan) at a frame rate of 12 frames per second (fps).

### 2.3 Light stimulation

For light stimulation experiments, four types of LEDs and three types of lasers were used (Supplementary Table S1). The white LED used for stimulation (LA-HDF158A, Hayashi Repic Co. Ltd., Tokyo, Japan) was identical to the illumination light source. While maintaining a constant imaging illumination intensity, an additional white LED light source (50,000 lx) was applied for stimulation. Under the same fixed illumination conditions, all other stimulation lights were directed to the targeted dermal region of the specimen. For RGB LED stimulation, blue, green, and red LEDs (MCEP-CB8-070-3, MCEP-CG8-070-3, and MCEP-CR8-070-3; Moritex Corp., Japan) were used, with peak wavelengths of 470 nm, 540 nm, and 630 nm, respectively. The illumination intensity of all LED stimulations was adjusted to 50,000 lx.

Blue laser stimulation (λ = 450 nm; FS-18001 A05-110 (450), Optronscience Inc., Japan) was applied at an intensity of 3,500 lx. Two high-intensity green lasers were also used: one at 520 nm with an intensity of up to 8,000 lx (FS-18002 A05-110 (520S), Optronscience Inc., Japan) and another at 530 nm with an intensity of 50,000 lx (Z-532-100T, Lucir, Japan). Detailed specifications of all stimulation light sources are summarized in Table S1. For both LED and laser stimulation, the optical axis of the emitted light beam was oriented perpendicular to the surface of the fish body.

### 2.4 Evaluation of light stimulation effects on dermal reflection

Initial qualitative assessments of the effects of strong LED or laser stimulation were performed using images captured by the CMOS camera through the mono-zoom lens. Quantitative analyses were conducted on real-time playback images displayed on a laptop computer (PG-GN1863YGF, LaVie, NEC, Japan), as described previously [26, 27].

## 3. Results and discussion

### 3.1 Photoresponse of Twinkling Iridophores to White LED Light Exposure

Atherinidae, including *Atherinomorus lacunosus* and *Hypoatherina tsurugae*, is a family of silversides. In previous studies [26, 27], a rapid twinkling and dynamic light reflection generated by guanine platelets in iridophores was first discovered in the iridophore membrane structure of the hardyhead silverside, *Atherinomorus lacunosus*. Similar twinkling behavior was also observed in the skin of the cobalt silverside, *Hypoatherina tsurugae*. Along the edges of the scales on the dorsal trunk of these fishes, membrane tissues composed of groups of iridophores (hereafter referred to as iridophore spots), approximately 100 µm in diameter, exhibited rapid and repetitive light reflection at frequencies of several hertz, independent of body motion.

The iridophore spots exhibited three distinct states: static bright, dynamically twinkling, and dark. During observations of the fish body surface, it was hypothesized that the initiation of twinkling was triggered by changes in illumination intensity. Therefore, the present study investigated the effects of controlled light stimulation in addition to the illumination used for observation. A white light-emitting diode (LED) was first employed as a potential stimulus. White light exposure was applied to a region containing several arrays of iridophore spots that were initially in a static bright state (Fig. 1A). The effect of stimulation was detected as a quenching of iridophore light reflection after termination of the white light exposure. Specifically, iridophore spots that were initially bright became dark immediately after cessation of the light stimulation, and subsequently recovered their brightness within a few seconds (Fig. 1B). White light stimulation was applied when the iridophore reflection was in the static bright state. The reflection intensity measured before stimulation was defined as the baseline. For quantitative analysis, the average reflection intensity measured over approximately four seconds prior to the additional light stimulation was normalized to 1.0. The effect of light stimulation was evaluated at two time points: immediately after termination of stimulation and during a four-second period following stimulation (Fig. 1C). In both evaluations, an average decrease of approximately 70% in reflection intensity was observed. The dependence of quenching on the exposure duration was further examined (Fig. 1D and Supplementary Fig. S2). The transition of the reflection state from bright to dark showed little dependence on the duration of light exposure (Fig. 1D). Notably, quenching of light reflection in the iridophores of Hypoatherina tsurugae was initiated within two seconds of additional white light stimulation.

**Fig. 1.**
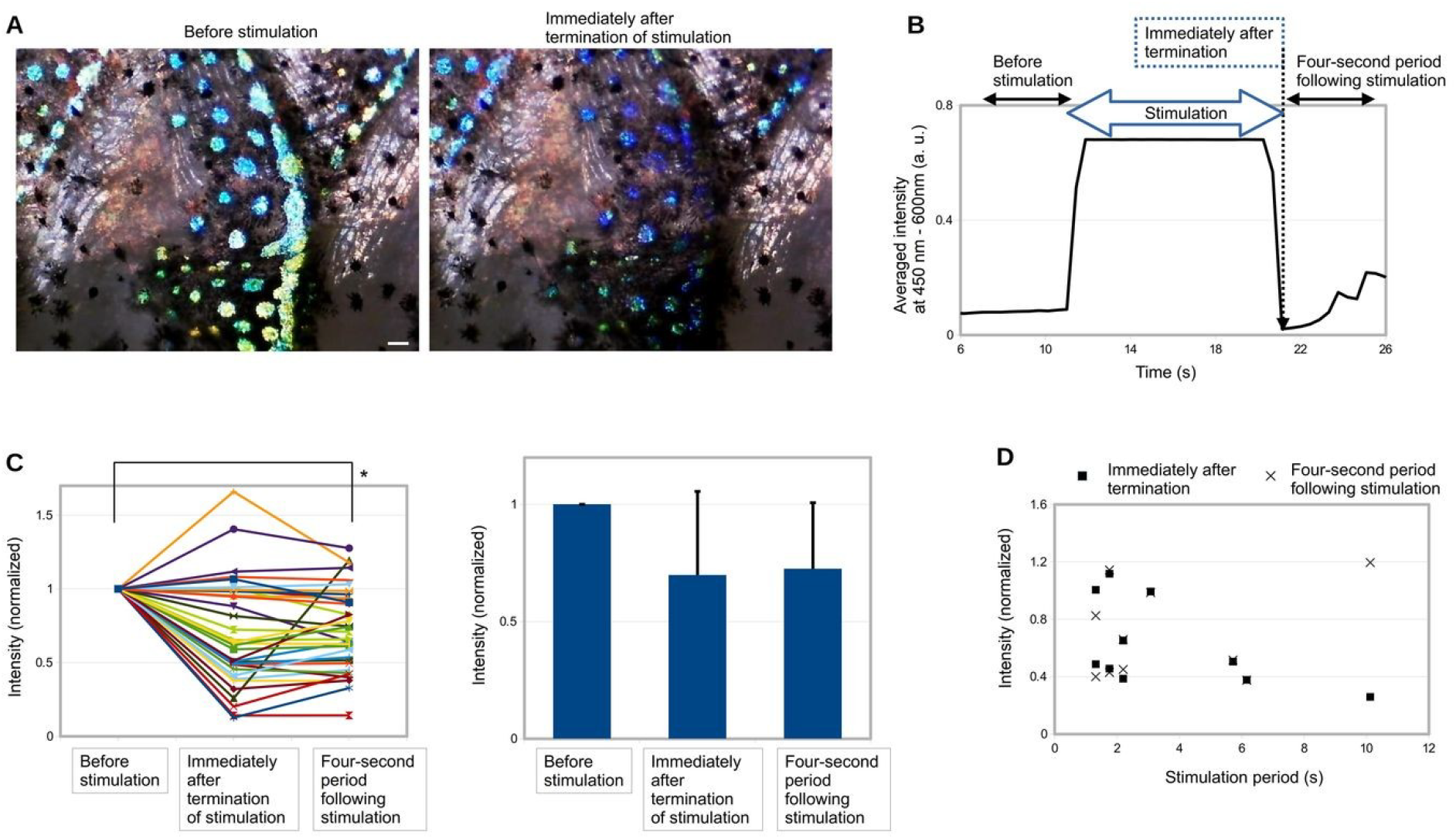
Effects of white LED stimulation on light reflection by iridophores in *Hypoatherina tsurugae*. (A) Quenching of light reflection from iridophores immediately after white LED stimulation. Scale bar, 100 µm. (B) Representative time course of reflected light intensity measured before, during, and after white LED stimulation. Optical intensity was quantified from recorded videos displayed on a monitor (monitor-based measurement). (C) Statistical analysis of quenching magnitude over time. Data for the pre-stimulation period represent the average over a four-second interval, as do data for the four-second interval immediately following stimulation. (D) Dependence of quenched light reflection intensity on the duration of white LED stimulation.

### 3.2 Comparison of Photoresponses to Different Colors of Light Stimulation

The possible contribution of individual light colors to the quenching of light reflection was investigated. First, the photoresponse of iridophore light reflection to blue illumination was examined (Fig. 2). The data shown in Fig. 2 were obtained using a blue laser (450 nm, 20 mW, 3,500 lx) as a coherent light source. Notably, iridophore spots aligned along the edges of the scales changed from a bright to a dark state upon blue light stimulation (Fig. 2A).

**Fig. 2.**
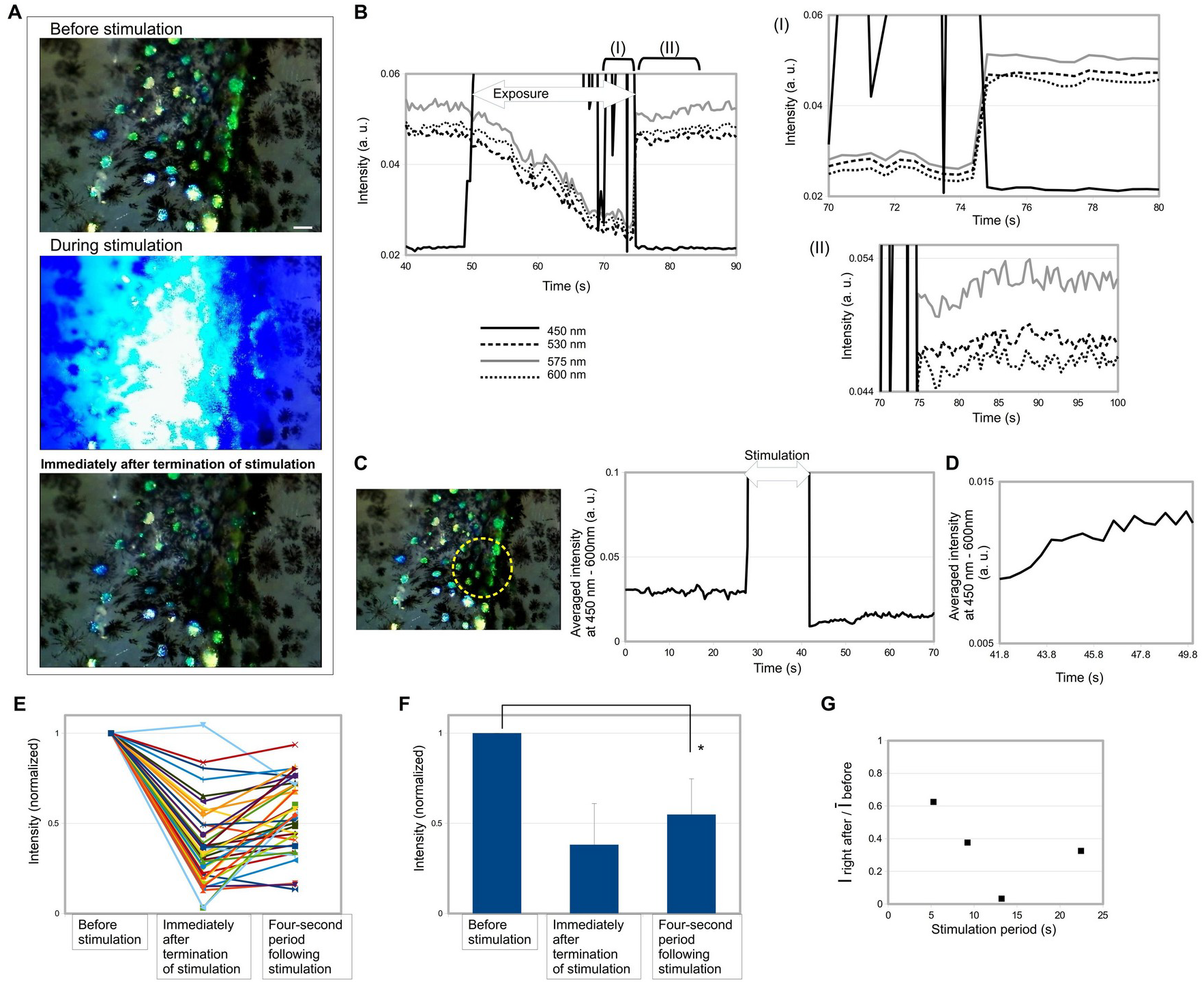
Effects of blue laser stimulation on light reflection by iridophores in Hypoatherina tsurugae. (A) Images of iridophore light reflection before, during, and immediately after blue laser stimulation. Scale bar, 100 µm. (B) Time course of reflected light intensity directly measured through the microscope objective using a spectrophotometer. Right panels show magnified views of the latter phase of stimulation. (C) Time course of quenching and recovery of light reflection in a selected region of interest, quantified by monitor-based measurement. (D) Magnified view of the post-stimulation period shown in (B). (E, F) Changes in reflected light intensity, averaged over four-second intervals before stimulation, immediately after stimulation, and during the four-second period following stimulation. (G) Dependence of quenching magnitude on the duration of blue laser stimulation.

Real-time spectroscopic measurements were performed in the region targeted for light stimulation (Fig. 2B). These measurements of the transient photoresponse process revealed that the photoresponse in the dermal tissue of Hypoatherina tsurugae began immediately after the onset of stimulation with the 450 nm laser. During laser irradiation, the light reflection intensities at three optical wavelengths (530 nm, 575 nm, and 600 nm) gradually decreased. The two right-hand graphs in Fig. 2B show magnified views of the time windows labeled periods I and II.

The magnified time-course graph for period I suggests that iridophore reflection intensity increased immediately when the laser irradiation was weakened. This rapid change in reflection is likely attributable to changes in the inclination of guanine platelets within the iridophores. Consistent with the observation of photoresponses occurring within two seconds (Fig. 1D), the blue laser irradiation experiment further indicates that initiation of the photoresponse occurs on a timescale of seconds.

During period II, the blue laser irradiation was completely terminated. The reflection intensity immediately after switching off the irradiation was lower than the baseline level, and subsequently recovered to the original baseline within approximately ten seconds. Another representative example (Fig. 2C) similarly demonstrates that the time constant for recovery from the dark to the bright state was approximately ten seconds. Statistical analyses (Fig. 2, E and F), comparing reflection intensities immediately after stimulation and during the four-second period following stimulation, confirmed the significance of the blue laser–induced effects. When the duration of light stimulation was varied from 5 s to 20 s, an exposure time of approximately 13 s was found to be most effective for inducing quenching (Fig. 2G, Supplementary Fig. S3, and S3).

Red light stimulation also induced quenching of iridophore reflections (Fig. 3). These experiments were conducted using a red LED at an illumination intensity of 50,000 lx. The reflection intensity decreased upon red light stimulation and subsequently exhibited a recovery process (Fig. 3A). The significance of this effect was confirmed by statistical analysis (Fig. 3B).

**Fig. 3.**
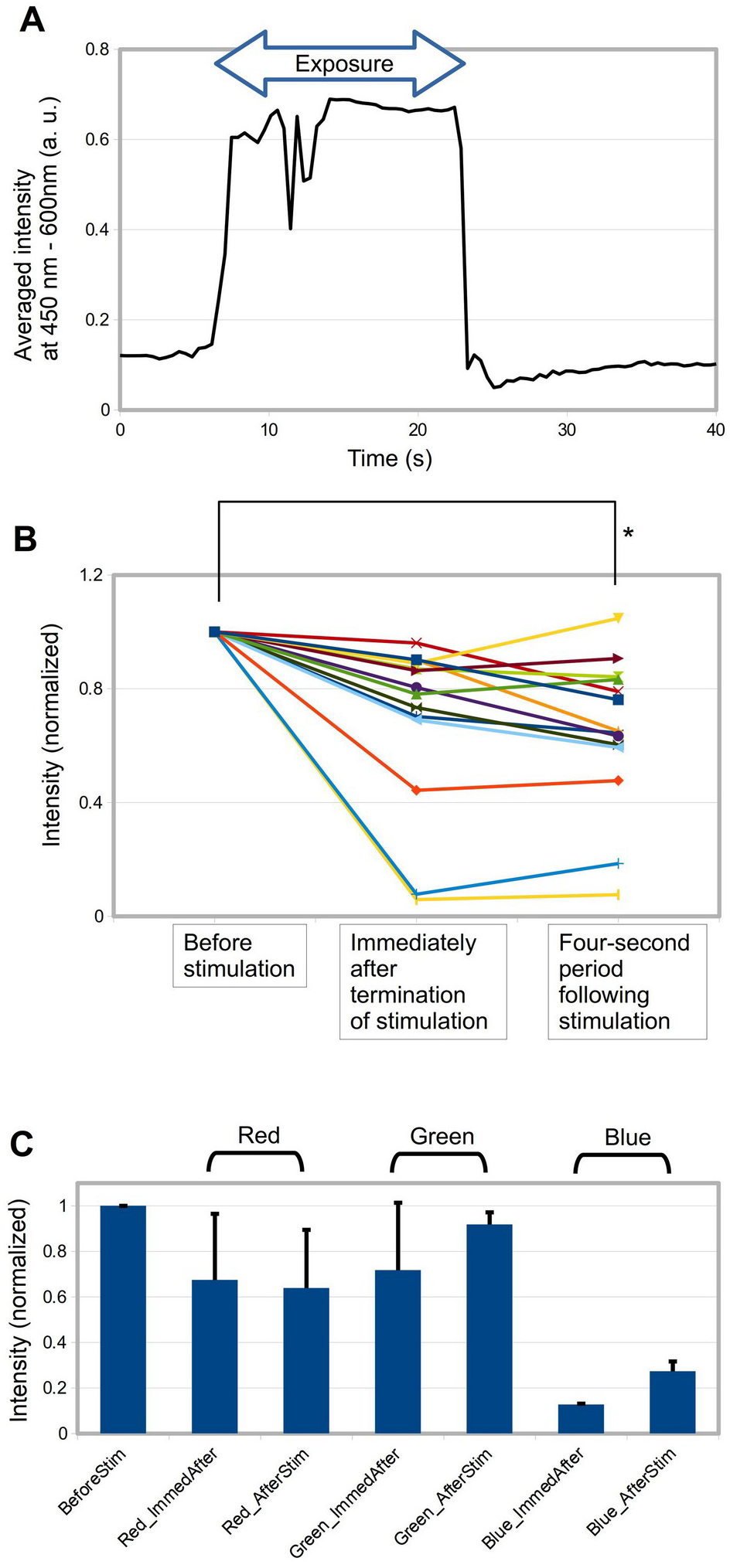
Quenching of light reflection in iridophore spots induced by red, green, and blue LED stimulation. (A) Representative time course of reflected light intensity following a single red LED pulse. (B) Comparison of reflected light intensity averaged over four-second intervals before stimulation, immediately after stimulation, and during the four-second period following red LED stimulation. (C) Comparison of quenching effects induced by red, green, and blue LED stimulation. Baseline intensity (BeforeStim) represents the four-second pre-stimulation average. Immediate post-stimulation intensities (Red_ImmedAfter, Green_ImmedAfter, Blue_ImmedAfter) and post-stimulation averages (Red_AfterStim, Green_AfterStim, Blue_AfterStim) are defined accordingly.

Using the same LED power source with interchangeable blue and green LED heads, the effects of individual light colors were directly compared (Fig. 3C). All irradiations were set to an intensity of 50,000 lx. A pronounced quenching effect, in which reflection intensity decreased to less than half of the baseline, was observed with blue LED light. In contrast, red and green light stimulation reduced reflection intensity by approximately 10–40%. Among the tested wavelengths, green LED illumination produced the weakest photoresponse.

The reduced effectiveness of green light stimulation was further examined using high-intensity green laser irradiation (Fig. 4). A notable feature of the green laser response was the rapid recovery of quenched iridophore reflection (Fig. 4A), with a recovery time constant shorter than those observed for blue and red light stimulation. Two possible interpretations can be proposed. One is a delay in the initial rapid realignment of light-reflecting guanine platelets, potentially involving a rotational process. The other is the involvement of an active control mechanism that accelerates the overall recovery process. Green laser irradiation at 8,000 lx produced a weaker quenching effect than blue laser irradiation, despite having approximately twice the illumination intensity (Figs. 4, B and C). Furthermore, increasing the green laser intensity to 50,000 lx did not enhance the quenching effect.

**Fig. 4.**
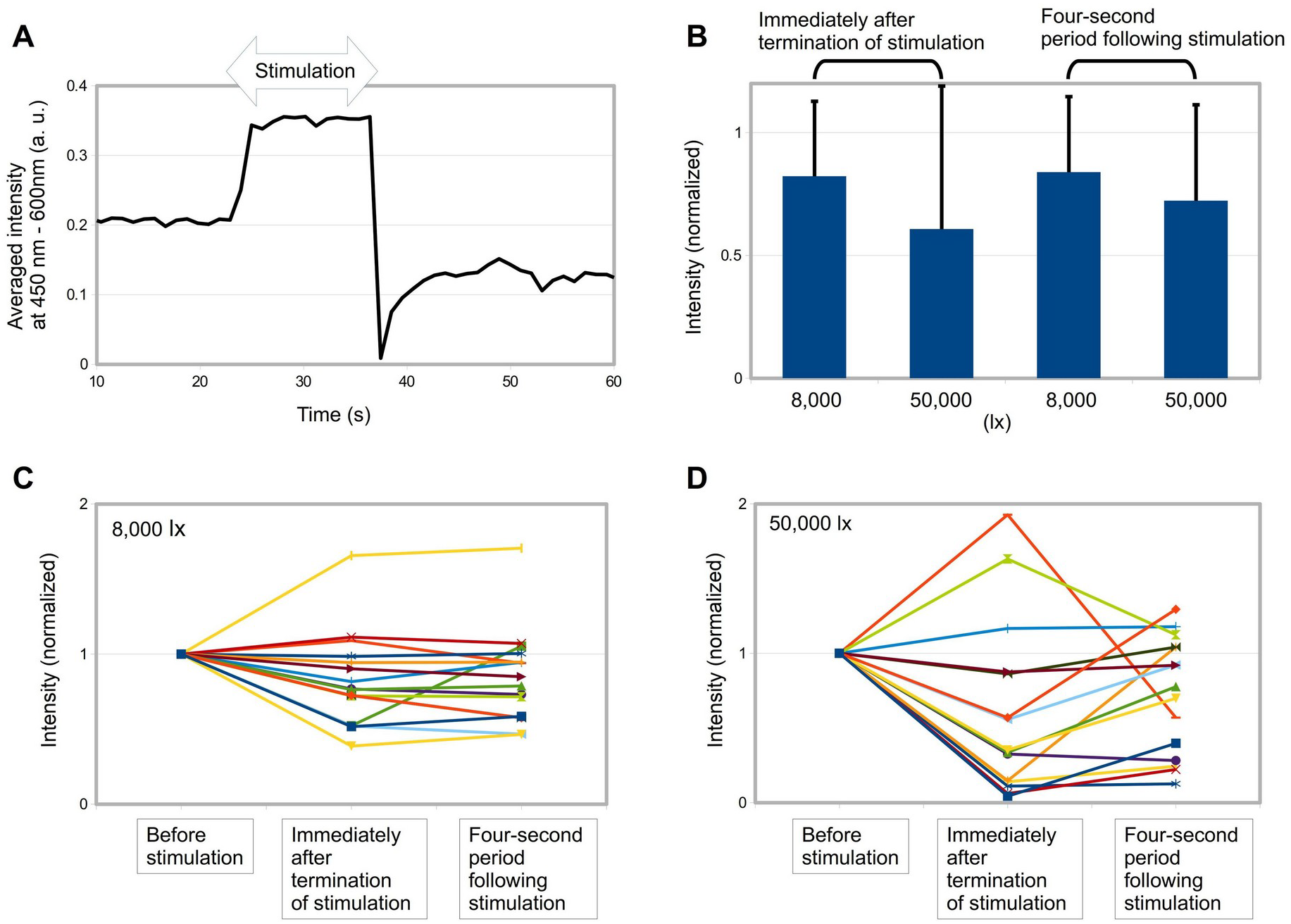
Quenching of light reflection in iridophore spots induced by green laser stimulation. (*A*) Representative time course following a single green laser pulse (530 nm, 50,000 lx). (*B*) Comparison of quenching effects induced by two classes of high-intensity green lasers. (*C*) Changes in reflected light intensity before stimulation, immediately after stimulation, and during the four-second post-stimulation period at 520 nm (8,000 lx). (*D*) Changes in reflected light intensity induced by stimulation at 530 nm (50,000 lx).

### 3.3 Discussion

The dominant colors of light reflected from the dermal tissue containing chromatophores of *Hypoatherina tsurugae* were yellow and blue (Supplementary Fig. S4). The light-reflecting chromatophore complex consists of iridophores, melanophores, and xanthophores. The yellow coloration is attributable to xanthophores. During light-induced quenching, the yellow coloration of the irradiated dermis disappeared simultaneously with the blue reflection. Previous studies have reported that aggregation of yellow pigment granules in xanthophores occurs within approximately 30 seconds; however, the transient and initial dynamics of pigment particle movement remain unclear. Because the present study focused on dermal light reflection, the contribution of xanthophore photoresponses to the observed quenching effect is likely minor compared with that of iridophores.

Several previous studies have reported that certain fish species exhibit photoresponses that are particularly sensitive to blue light. For example, aggregation of yellow pigment granules in xanthophores was observed in medaka under strong blue light (∼8,000 lx) [18]. Blue-light–dependent pigment movement in octopus skin has also been detected under illumination intensities of up to 150 µmol m^−2^ s^−1^ [2, 8]. Studies on octopus photoresponses have further demonstrated the presence of opsin molecules responsible for blue-light sensing. In the present experiments, the blue laser provided photon fluxes of approximately 1,000 µmol m^−2^ s^−1^ at an illumination intensity of 1,000 lx. Although this study did not include molecular analyses of opsins in the photoresponsive tissue, it does not exclude the possibility that opsin molecules in the skin of *H. tsurugae* function as optical sensors.

A major strength of this study lies in its ability to capture transient photoresponses with sub-second temporal resolution in a teleost species. The timescale of the observed quenching dynamics differs markedly from that reported in many previous studies, which focused primarily on the relatively slow aggregation or dispersion of chromatophore pigment particles. A recent report on octopus arm photoresponses demonstrated neural action potentials with millisecond temporal resolution [2]. Notably, the temporal dynamics of twinkling iridophore spots in *H. tsurugae occur* on a similar timescale to neural activity. Therefore, the switching between bright and dark iridophore states may provide information about underlying neural activity in response to light stimulation. Furthermore, it may be possible to analyze spatiotemporal patterns of neural networks controlling both twinkling behavior and optical sensing by chromatophore complexes in dermal tissue. In this context, *H. tsurugae* represents a promising model for exploring neuronal innervation of skin-embedded photoresponsive systems.

Observations of phototactic behavior in aquatic organisms, such as photosynthetic microorganisms [28, 29] and nematodes (*Caenorhabditis elegans*) [30], have provided insights into biological light-sensing mechanisms. Currently, opsins are considered primary candidates for the core components of biological photoreception across diverse taxa. The rapid optical sensing observed in *H. tsurugae* likely involves such common mechanisms, including opsins or photosensitive adenylate cyclases, particularly for blue-light detection. However, the measurable responses to strong green light irradiation suggest the presence of an additional optical sensing mechanism distinct from that mediating blue-light responses.

Possible mechanisms underlying non-ocular, skin-embedded biological optical sensing can be broadly categorized into thermal and non-thermal processes. Non-thermal mechanisms include photoreception mediated by biomolecules, such as opsins, that convert photon energy into neural signals. In contrast, thermal effects of light irradiation include photothermal conversion [31, 32], high-density plasma generation and optical breakdown [33, 34], and ablation [35–40]. In the present study, no ablation was observed in light-irradiated tissues. Any photothermal effects induced are therefore expected to be weak and likely operate at non-thermal or quasi-non-thermal levels. If photothermal energy conversion generates localized mechanical forces, such forces could induce local tissue deformation and stimulate nearby neurons. This type of sensor could coexist with photosensitive molecular sensors, such as opsins, rhodopsins [29], and photoactivated adenylate cyclases [28]. Neural networks receiving signals from photosensitive molecules may also integrate inputs from neighboring sensors operating via physical mechanisms. In addition to photothermal effects, photo-orientation [41, 42] and radiation pressure [43, 44] may induce movements of micro- and nano-scale structures, including molecules and organelles, within skin tissues.

Complexes of molecules or cells capable of absorbing photon energy may serve as transducers that relay optical signals to the nervous system. The optical sensory system of *H. tsurugae* may comprise modular components that respond preferentially to strong versus moderate sunlight intensities. To further elucidate fish photoresponses, it is useful to hypothesize that multiple sensory modalities—including thermal and non-thermal mechanisms, as well as cross-modal interactions—operate simultaneously. The present study revealed stochastic photoresponses to green light stimulation, highlighting the importance of detailed, time-resolved analyses for achieving a deeper understanding of skin-embedded optical sensor systems.

Recently, the dermis and epidermis of human skin have inspired advances in wearable electronics and skin-mounted sensor devices [45]. Wearable technologies aim to enable stretchable, lightweight, and ubiquitous health-monitoring systems [46–49], often designed to interface intimately with living skin tissue. Investigating the skin structures of diverse animal species may therefore provide novel approaches to skin-device innovation. Existing wearable devices support applications including healthcare monitoring [46], pressure sensing [47], epidermal molecular transport detection [48], and haptic signal processing [49]. Moreover, bio-inspired devices capable of infrared-reflective camouflage [50] demonstrate the high potential of biomimicry for developing advanced epidermal technologies. From a biomimetic perspective, fish represent an attractive comparative model for humans. Mimicking newly discovered optical functions of fish skin may contribute to the development of wearable devices optimized for operation under strong sunlight.

## 4. Conclusion

In conclusion, the dermal tissue of H. tsurugae possesses an optical sensing system that responds most strongly to blue light at 450 nm. This photoresponse is characterized by the disappearance of light reflection from dermal iridophores. Similar but weaker effects were observed under green and red light stimulation. The reduced efficacy of green and red light, together with differences in response dynamics, suggests that distinct sensory mechanisms are involved depending on the wavelength of light stimulation.

Masakazu Iwasaka: Investigation, Methodology, Experimental design, Experiments, Analysis, Data collection Conceptualization, Writing.

## Funding

No funding

## Declaration of competing interest

The authors declare that they have no known competing financial interests or personal relationships that could have appeared to influence the work reported in this paper.

## Acknowledgments

Author appreciates the university institute for providing the experimental space.

## Data availability

All data and materials used in the analysis are available in the main text, the supplementary materials or Zenodo repository.

## Notes

### Competing Interest Statement

The authors have declared no competing interest.

