## Supplementary Material for "Ultrafast light sensing mediated by dermal iridophores in the silverside fish *Hypoatherina tsurugae*"

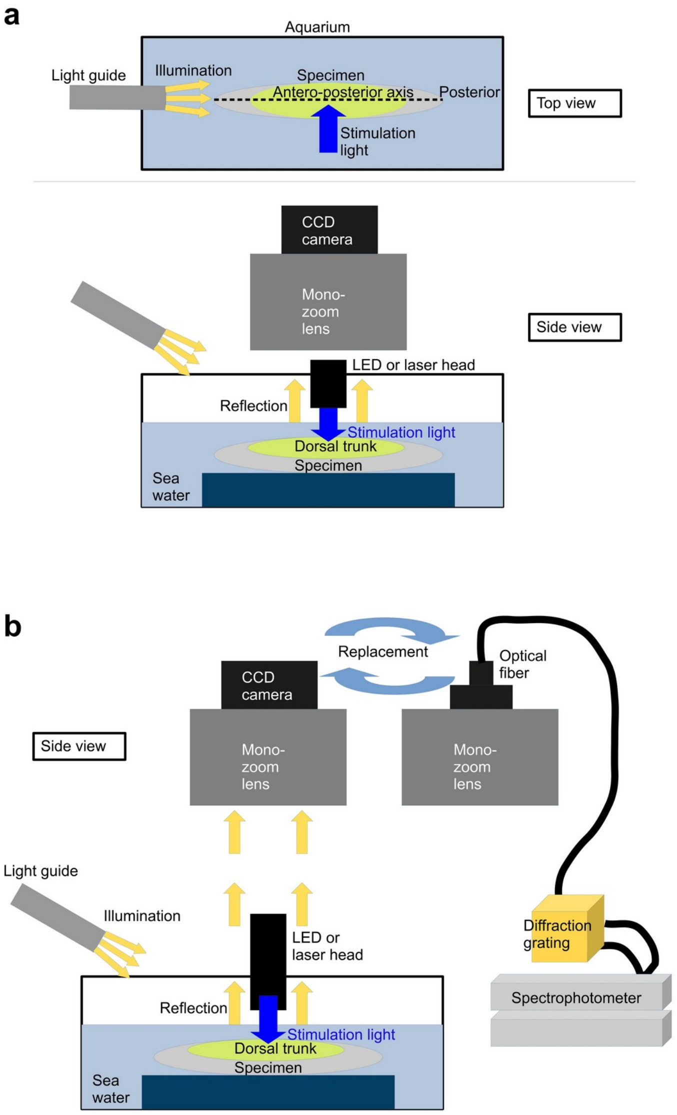


Fig. S1. Experimental setup for observing light-induced effects on twinkling spots on the dorsal trunk of *Hypoatherina tsurugae*. (a) Top and side views of the experimental setup, including the aquarium, observation illumination, zoom lens, CMOS camera, and stimulation light source. The top view illustrates the spatial alignment of illumination beams and specimen orientation. (b) Switching between microscopic imaging and visible-light spectroscopy by replacing the CMOS camera with an optical fiber connected to a spectrometer.

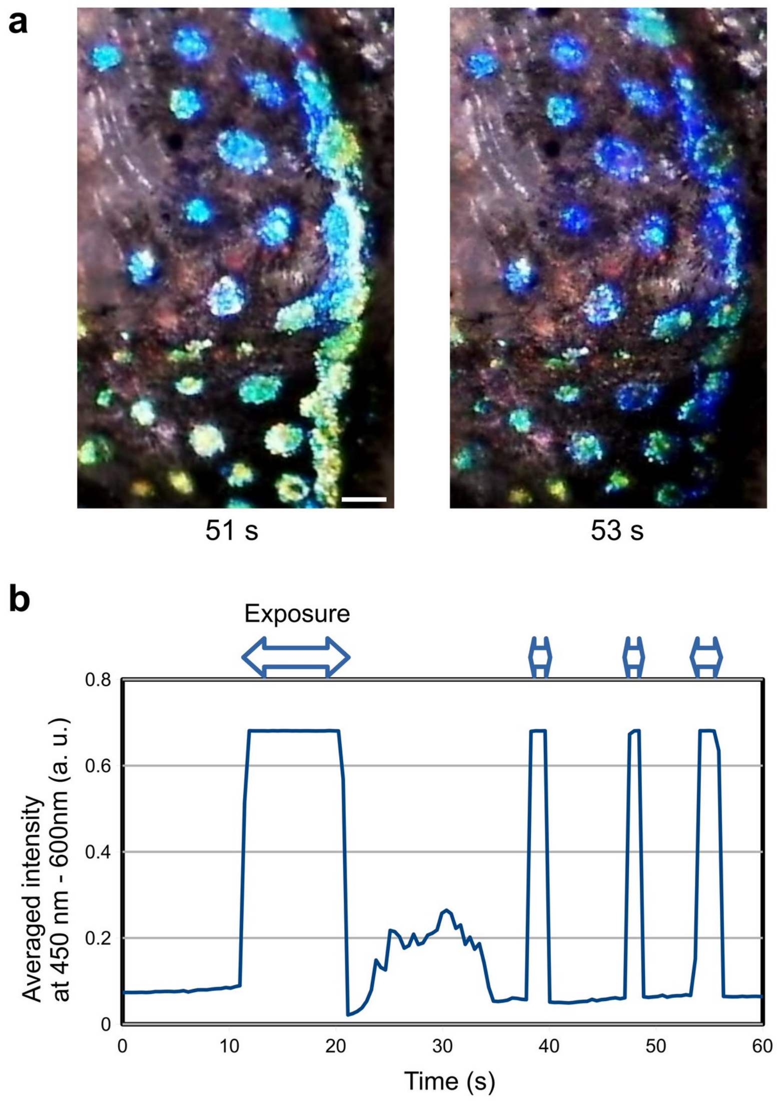


Fig. S2. Quenching of light reflection in twinkling spots of *H. tsurugae* induced by white LED stimulation. (a) Representative images before (left) and after (right) stimulation. Time stamps correspond to the time points shown in (b) and Supporting Movie 1. (b) Time course of averaged dermal reflected light intensity during four repeated white LED stimulations.

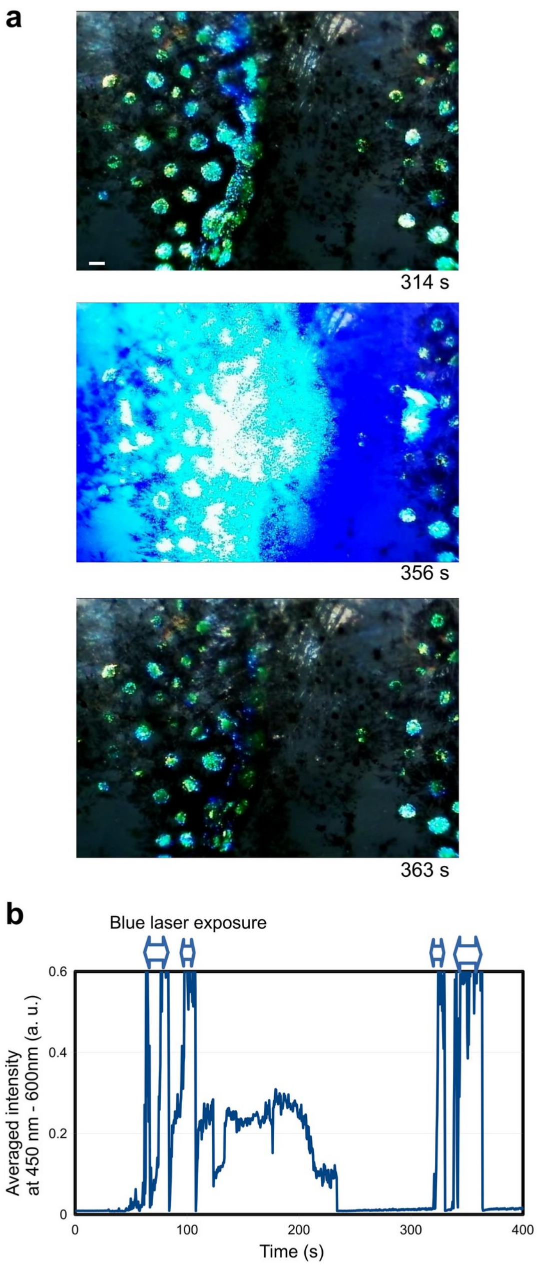


Fig. S3. Quenching of light reflection in twinkling spots of *H. tsurugae* induced by blue laser stimulation. (a) Representative images before (top), during (middle), and after (bottom) stimulation. Time stamps correspond to (b). (b) Time course of averaged dermal reflected light intensity during four repeated blue laser stimulations.

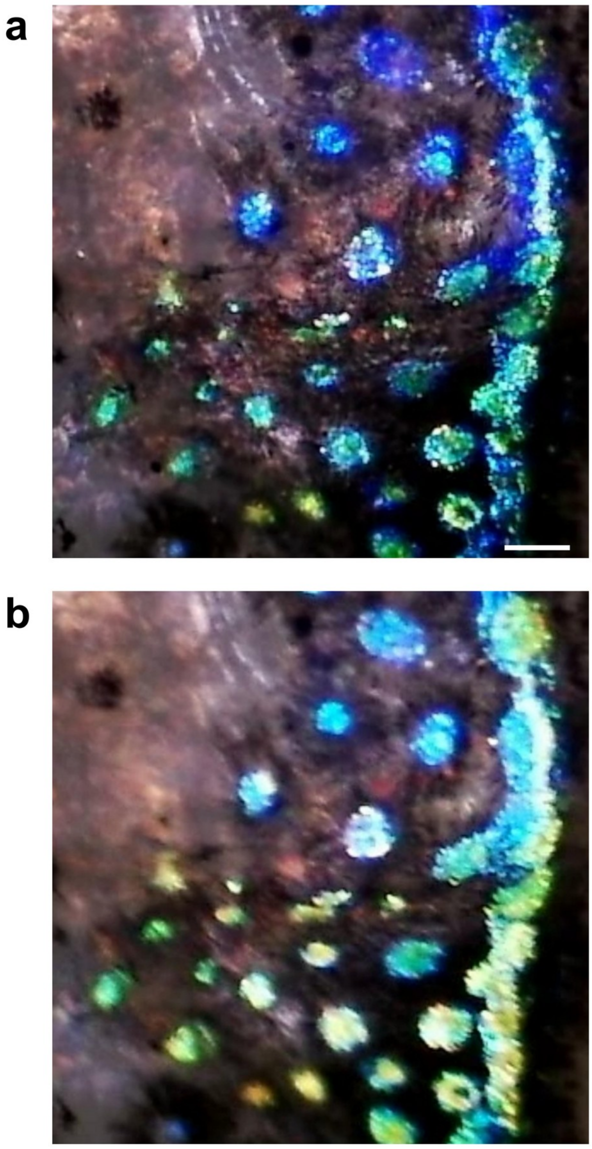


Fig. S4. Two representative reflected-light color states in *H. tsurugae*. (a) Blue-dominant state. (b) Yellow-enhanced state.

Table S1. Specifications of light sources used for stimulation.

| Type | Color (peak wavelength (nm)) | Illumination intensity (lx) | Maker & specification |
|---|---|---|---|
| LED | White (450, 550) | 50,000 | LA-HDF158A, Hayashi Repic Co. Ltd, Tokyo, Japan |
| LED | Blue (470) | 50,000 | Moritex Japan |
| LED | Green (540) | 50,000 | Moritex Japan |
| LED | Red (630) | 50,000 | Moritex Japan |
| Laser | Blue (450) | 3,500 | Optronscience, Japan |
| Laser | Green (520) | 8,000 | Optronscience, Japan |
| Laser | Green (530) | 50,000 | Lucir, Japan |
